# Functional Heterogeneity of Voice-Encoding Cortex Revealed by Clinical Language Mapping

**DOI:** 10.1101/2024.05.24.595794

**Authors:** Emily E. Harford, Kyle M. Rupp, William P. Welch, Ruba Al-Ramadhani, Avniel Ghuman, Lori L. Holt, Taylor J. Abel

## Abstract

Regions in the superior temporal sulcus and gyrus have been heavily implicated in voice-selective responses in human auditory cortex. Despite an apparent specialization for the encoding of human voice, research outside the auditory domain suggests that these areas likely participate in additional neural processes including speech processing and production. The aim of the current study was to combine results of electrophysiological recording and clinical stimulation mapping procedures in patients undergoing stereoelectroencephalography (sEEG) to explore potential functional heterogeneity in voice-encoding cortex. Both channels that demonstrated voice-encoding properties and channels critically implicated in language functioning were heavily concentrated in the left STG/S. Analysis of functional overlap revealed channels in the posterior STG/S that appear to be involved in both voice encoding and language. Strength of voice encoding in these functionally diverse sites was not significantly different from sites that were implicated in voice encoding alone. Our findings add to prior observations of functional heterogeneity in the STG/S and contribute to proposed models of speech perception. We discuss these results in the context of the utility of electrophysiological methods in mapping cortical networks and identifying regions essential for functioning.

## 1 Introduction

Research spanning decades has sought to explain the neural basis of conspecific voice selectivity, or the observation that own-species vocalizations elicit stronger neural responses in certain regions of the auditory cortex than other auditory categories. Vocalizations relay crucial information to listeners about the identity, emotions, and other characteristics of the source (see Schweinberger et al., 2014; Seyfarth & Cheney, 2003 for reviews), indicating an ecological and social significance for maintaining a privileged status for conspecific voice processing. In human auditory cortex, these regions are typically referred to as “temporal voice areas” (TVAs) and are localized in the bilateral upper bank of the superior temporal sulcus (STS) and the anterior superior temporal gyrus (STG) (Agus et al., 2017; Belin et al., 2000, 2002; Bodin et al., 2018; Fecteau et al., 2004; Pernet et al., 2015).

Studies in auditory neuroscience focus on these regions as highly specialized for processing human voice, while research outside the auditory domain suggests that the STS is functionally diverse (Hein & Knight, 2008; Liebenthal et al., 2014). Regions identified as TVAs in the voice processing literature are also implicated in audiovisual (Stevenson & James, 2009; Watson et al., 2014), biological motion (Beauchamp et al., 2002; Grossman et al., 2005), face (Haxby et al., 2000; Winston et al., 2004), and speech (Binder et al., 2000; Redcay, 2008) processing.

Though functional specialization is a core principle of cortical organization, it is likely that some overlap exists. While both lesion and transcranial magnetic stimulation studies suggest a causal role for TVAs in voice perception, other lines of evidence suggest a diversity of other functions.

In this manuscript, we explored potential functional heterogeneity in voice-encoding regions of human auditory cortex. We capitalized on the unique opportunity to combine electrophysiological responses during auditory stimulus presentations with observations from cortical stimulation mapping to investigate whether regions involved in voice encoding may also play a role in language functioning.

## 2 Methods

### 2.1 Participants

Participants selected for this study were neurosurgical patients with drug-resistant epilepsy undergoing stereoelectroencephalography (sEEG) monitoring as part of staged evaluation for epilepsy surgery. Patients were included if they completed a target auditory task and underwent clinical stimulation mapping (discussed further below in *2.2* and *2.3*) (n=14). Furthermore, patients were excluded if clinical stimulation mapping procedures did not produce a clinical effect on speech/language (n=7) or if disruption to speech/language was thought to be related to motor speech/articulatory control (i.e., clinical effect upon stimulation of channels in the pre- or motor cortex) (n=3). These inclusion and exclusion criteria resulted in a final sample of 5 participants (mean age=16.2 years).

### 2.2 Neural Data Collection and Auditory Task Procedures

Neural data were obtained during sEEG recording as participants completed an auditory 1-back task. This task, which we refer to as Natural Sounds (NatS) (Norman-Haignere et al., 2015), has been used in previous research (Norman-Haignere et al., 2022; Rupp et al., 2022) to determine voice-encoding channels. The NatS stimuli consist of 165 sounds, each 2 seconds in duration, belonging to 1 of 11 auditory categories. These categories include human voice (both speech and non-speech vocalizations), musical instruments, sounds from nature (e.g., running water), animal vocalizations, and other environmental sounds (e.g., car horn). Stimuli were delivered binaurally through Etymotic ER-3C insert earphones with volume set to a comfortable level for each patient during an audio calibration step performed prior to the beginning of the experiment. Participants provided responses to 1-back repetition of stimuli by pressing a button (Response Time Box, v6). Neural data were recorded at 1kHz via a Ripple Grapevine Nomad processor with notch filters applied at 60, 120, and 180 Hz.

### 2.3 Clinical Stimulation Mapping Procedures

Cortical stimulation mapping was performed for each participant as dictated by clinical utility to localize eloquent cortex. Stimulation was delivered at a range of 2-10mA, a pulse rate of 50Hz, a pulse width of 200-250 microseconds, and a duration of 1 second with the goal of producing a clinical effect on motor or speech production. Tasks included counting, reading aloud, generative naming, confrontation naming, and spontaneous speech, and were varied across patients according to developmental level at the discretion of the clinical team (Table 1). Bipolar current was delivered between two contiguous electrode contacts as patients performed these tasks.

**Table 1.**
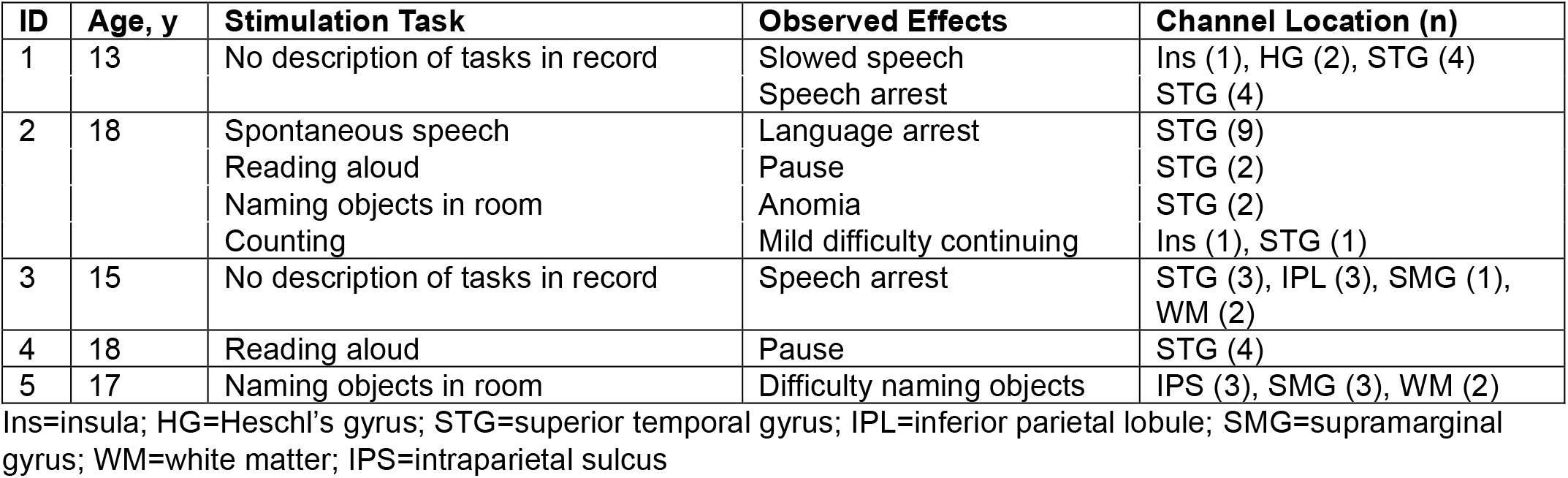
Participant, stimulation task and stimulation effect details by location of stimulation applied.

The clinical team documented any clinical effect observed in the medical record.

### 2.4 Identification and Functional Categorization of Channels

To examine potential overlap between regions in the voice processing and language networks, we labeled channels according to their apparent functional involvement in voice encoding and language functioning. Channels were divided into three categories: channels that showed a category preference for human voice over other sound categories (VOICE+), channels that produced a disturbance in language functioning when stimulated (STIM+), and channels that met both these criteria (OVERLAP). VOICE+ channels were identified based on neural data recorded during the auditory NatS task while STIM+ channels were identified based on documentation from clinical stimulation mapping procedures.

To identify VOICE+ channels, we followed the procedure from Rupp et al. (2022) and selected those channels that not only showed greater high gamma activity (HGA) responses to voice over nonvoice sounds but also appeared to encode voice at the categorical level. Briefly, HGA was calculated by filtering common-average-referenced data between 70-150 Hz and using the Hilbert transform to extract the analytic amplitude of the filtered signal. All channels were subjected to a nested encoding model analysis that attempted to predict HGA responses (averaged across a window of 480-1450 ms post-stimulus onset) using stimulus features estimated across this same window. This model considers both lower-level acoustic properties of stimuli and auditory category, accounting for inherent variability in the acoustic profiles of sound categories that may drive neural responses by incorporating a binary category feature for human voice. A nested model that includes only acoustic features was compared via a likelihood ratio test to a full model that includes both acoustics and the binary category variable. In calculating the likelihood ratio test statistic (χ^2^ distributed) between the model with and without the voice category feature, we estimate the likelihood that the category itself explains neural responses beyond low-level acoustic features. Channels were labeled as VOICE+ if the addition of the category feature resulted in a statistically significant improvement in the goodness-of-fit for the encoding model (p < 0.01, false discovery rate corrected). Relative strength of voice category encoding is represented by χ^2^ values, with larger values indicating an increased voice-encoding strength. See Rupp et al. (2022) for more details. To ensure that we accurately represented the degree of functional overlap, we included only those VOICE+ channels that were stimulated during clinical stimulation mapping procedures. Because all participants demonstrated typical left hemisphere lateralization in language function, stimulation was performed in only left hemisphere channels in 4/5 participants. Therefore, our analysis of voice-encoding channels is constrained to the left hemisphere.

STIM+ channels were identified by reviewing documentation of clinical stimulation mapping procedures in the medical record. Channels were labeled as STIM+ if they produced disturbance in performance of language tasks upon stimulation. Disturbance was defined as any change in the quality or efficiency of speech or language production and included decrease in speech quality (i.e., slurring or slowing of speech), naming difficulty, total speech arrest, and any other decreases in performance relative to performance prior to delivery of stimulation. As current was delivered between contacts themselves, both contacts in the pair were labeled as STIM+. As mentioned previously, we excluded channels in premotor, supplementary motor, and primary motor cortex that were believed to have produced a disturbance in task performance that was oral motor in nature.

Any channels that were labeled as both VOICE+ and STIM+ were then classified as OVERLAP channels. These channels both demonstrated evidence of categorical encoding of human voice and resulted in disturbance to speech/language function upon stimulation.

### 2.5 Channel Localization

Cortical surfaces were reconstructed for each patient using Freesurfer (Fischl et al., 2004) and a pre-operative MRI study. The MRI was co-registered with a post-operative CT scan using the Brainstorm package on MATLAB (Tadel et al., 2011). Next, MNI normalization was performed via a Brainstorm implementation of SPM12’s nonlinear warping algorithm. The MNI field was used to project the Julich atlas onto patient anatomy (Amunts et al., 2020; Eickhoff et al., 2005). Channels were localized to an atlas region-of-interest label according to the label of the closest voxel. These labels were visually inspected and corrected as needed.

## 3 Results

### 3.1 Regions Implicated in the Voice-Encoding Network (VOICE+ channels)

After excluding VOICE+ channels that were not stimulated during functional mapping, results of the encoding model analyses revealed an average of 12.4 channels (range 3-21) per patient exhibiting voice category preference. VOICE+ channels were largely concentrated throughout the STG/STS (n=48, 77.4%). Additional channels demonstrating voice-encoding properties were also observed in Heschl’s gyrus (HG), supramarginal gyrus (SMG), middle temporal gyrus (MTG), and insula (Table 2). Using a Wilcoxon rank-sum test (α = 0.05), we compared the relative strength of voice encoding between primary auditory (HG) and auditory association (STG/STS) regions. Results showed evidence of stronger voice category preference in channels located in STG/STS versus those located in HG (p=0.02).

**Table 2.** Distribution of VOICE+, STIM+, and OVERLAP channels by ROI label.

| Location | VOICE+, n chans | STIM+, n chans | OVERLAP, n chans |
| --- | --- | --- | --- |
| STG/STS | 48 | 24 | 20 |
| HG | 10 | 2 | 2 |
| SMG | 2 | 4 | 1 |
| MTG | 1 | 0 | 0 |
| Insula | 1 | 2 | 0 |
| IPL | 0 | 3 | 1 |
| WM | 0 | 4 | 0 |
| IPS | 0 | 3 | 0 |

STG=superior temporal gyrus; STS=superior temporal sulcus; HG=Heschl's gyrus; SMG=supramarginal gyrus; MTG=middle temporal gyrus; IPL=inferior parietal lobule; WM=white matter; IPS=intraparietal sulcus

### 3.2 Regions Implicated in Language Functioning (STIM+ Channels)

A range of 20-59 (mean 33.4) contact pairs were stimulated per patient, which included stimulation to induce clinical effects on both speech and motor functioning. For the purposes of this paper, we focused solely on those channels which produced a disturbance in speech and language tasks. Stimulation of an average of 8.4 channels (range 4-11) per patient produced a clinical effect during these tasks and were thus labeled as STIM+ channels. A majority of these channels were also concentrated in the STG/STS (n=24, 57.1%), with additional channels localized to the intraparietal sulcus (IPS), inferior parietal lobule (IPL), SMG, insula, and HG. Several channels (n=4, 9.5%) were situated in white matter and did not correspond to an atlas label.

### 3.3 Overlapping and Dissociated Regions (OVERLAP Channels)

To examine functional heterogeneity within voice-encoding regions, we identified those channels which were labeled as both VOICE+ and STIM+, which were then labeled as OVERLAP. The mean number of OVERLAP channels per patient was 4.8 (range 0-9), with only 1 patient demonstrating no common sites. OVERLAP channels were localized to the STG/STS (n=20, 83.3%), and HG (n=2, 8.3%), with single channels observed in the SMG and IPL.

Examination of the anatomical distribution of channel categories reveals evidence of both functional overlap and functional division, specifically within the STG/STS (Figure 1). Channels that play a critical role in language functioning but do not demonstrate properties of voice category encoding (STIM+ only) were observed in extreme posterior regions of the STG/STS as well as extratemporal regions including the supramarginal gyrus and intraparietal sulcus. In contrast, channels in middle/anterior STG/STS were almost exclusively those that demonstrated voice encoding properties but no disturbance of speech/language functioning during stimulation (VOICE+ only). Functional overlap in voice encoding and speech/language function was localized primarily to posterior regions of the STG/STS. A Wilcoxon rank-sum test (α = 0.05) comparing relative strength of voice encoding between VOICE+ only (i.e., VOICE+ with no effect on speech/language during stimulation) and OVERLAP channels found no significant differences (p=0.39).

**Figure 1.**
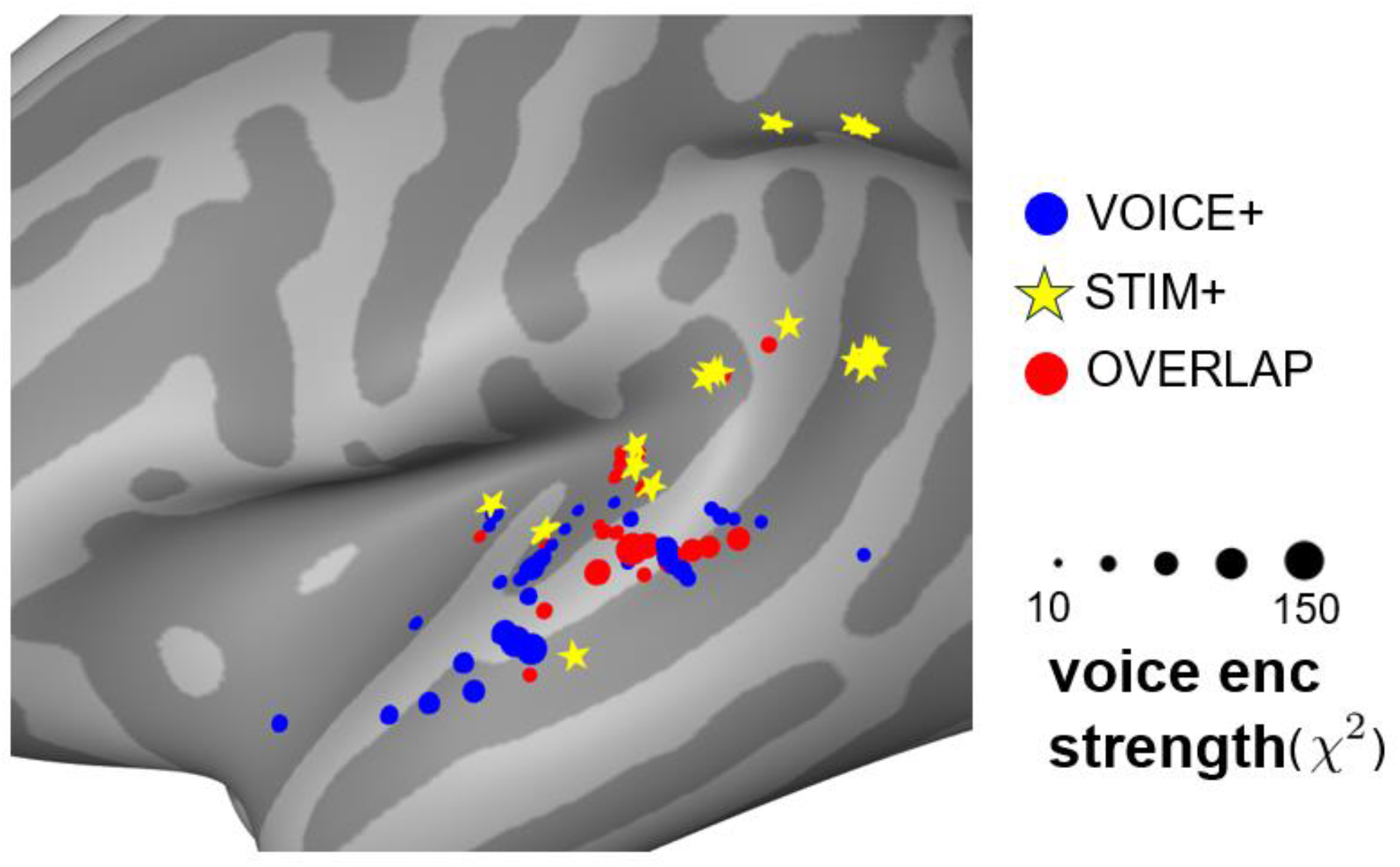
Distribution of channels implicated in language functioning only (STIM+), voice-encoding only (VOICE+), and in both (OVERLAP) including strength of voice encoding (voice enc strength) for VOICE+ and OVERLAP channels.

## 4 Discussion

The results of this exploratory study provide a unique perspective on functional diversity within voice-encoding regions in the human temporal lobe. Our primary aim was to examine potential involvement of voice-encoding regions in additional cognitive processes. In combining observations from clinical stimulation language mapping and electrophysiological recordings during an auditory task, we demonstrate that regions along the STG/STS appear to participate in both voice-encoding and language functioning. Our analysis identified three clusters illustrating functional dissociation and overlap: the middle/anterior STG/STS showed voice encoding but not language functioning, extreme posterior regions of the STG/STS extending toward the parietal lobe showed language functioning but not voice encoding, and the posterior STG/STS demonstrated evidence for both.

Our findings of functional overlap between voice-encoding and language functioning in the STG/STS fit well with prior research identifying both voice-selectivity (Agus et al., 2017; Belin et al., 2000; Pernet et al., 2015) and language functioning (Haglund et al., 1994; Hamberger et al., 2001; Kabakoff et al., 2024; G. Ojemann et al., 1989; G. A. Ojemann, 1991; G. Ojemann & Mateer, 1979) throughout this region. While studies in the voice processing literature tend to view TVAs under a narrow lens, it is clear that these regions participate in a wider variety of functions. In 4/5 of our participants, several voice-encoding channels also appeared to play a critical role in language functioning as evidenced by task disruption during clinical stimulation mapping. The existence of these OVERLAP channels suggests that putative TVAs are not strictly devoted to voice encoding; in other words, selectivity does not necessarily imply functional exclusivity. The observation that OVERLAP channels were primarily confined to the left posterior STG/STS is in agreement with a meta-analysis by Liebenthal et al. (2014), who identified the left posterior STS as a particularly versatile region within STS that was activated by the largest number of functional task categories across studies. Furthermore, the fact that there was not a significant difference in the strength of voice encoding in strictly VOICE+ channels versus OVERLAP channels indicates that functional diversity does not come at the cost of sensitivity in the case of voice encoding.

While several STIM+ channels were identified outside of the temporal lobe in our participants, over half (57.1%) were localized to the STG/STS. This finding suggests that auditory association cortex is implicated not only in perception, but also production of speech and language. This idea is well-substantiated by prior findings of left posterior STG/STS involvement in both speech perception and production tasks (Buchsbaum et al., 2001; Ekert et al., 2021; Hamberger, 2007; Hickok et al., 2009; Okada & Hickok, 2006; Wise et al., 2001). The localization of STIM+ only channels (i.e., those that produced a clinical effect on language but did not demonstrate voice-encoding properties) agrees with models of speech processing that propose left pSTS as a hub for auditory-motor integration (Hickok & Poeppel, 2007). Several groups have proposed that the left pSTS may be involved in speech production by maintaining short-term auditory/acoustic representations that are communicated and converted to articulatory representations in the motor network (Hickok et al., 2000, 2009; Wise et al., 2001).

One important caveat to the interpretation of these findings is that the two methods employed to categorize the function of channels, i.e., stimulation mapping and electrophysiological recording, may identify two different types of cortical regions that comprise a network: those that are critical to a function and those that contribute. The distinction of mapping “essential” areas via cortical stimulation versus “participating” areas via neural imaging or recording was first highlighted by Ojemann et al. (1989), who concluded that maps of participatory regions for a particular cognitive function are often more widely distributed than maps of essential regions. Our results illustrate this difference through the finding that cortical stimulation did not reveal a critical role of the anterior STG/STS in language production despite evidence that this region actively participates in speech and language processing and production (Farias et al., 2005; Friederici et al., 2000, 2003; Visser & Lambon Ralph, 2011). The results of Hamberger et al. (2001), which identified the anterior STG as critical to performance of auditory but not visual naming tasks, emphasize that identification of these essential regions is sensitive to task selection. On the other hand, electrophysiological recording and encoding model analysis revealed a voice-encoding network distributed throughout the STG/STS. Identification of regions essential to voice encoding or auditory category encoding would require the use of stimulation during an appropriately selected task (e.g., auditory categorization task) to observe the nature and localization of disruption in this process.

Studies identifying a lack of cohesion between language areas identified via fMRI or sEEG versus those identified via cortical stimulation mapping underscore this issue (Corina et al., 2000; Cuisenier et al., 2020; Roux et al., 2003). While identification of critical regions can be achieved through cortical stimulation and is considered standard clinical practice, neuroimaging and electrophysiological techniques are required to reveal the spatiotemporal dynamics of cognitive processes and the networks that subserve them. Though the latter is not substantially specific to guide surgical decision making, it is crucial in informing our understanding of functional division and specialization in the human brain. The marriage of these techniques in future research provides the best opportunity to describe the localization, mechanisms, and spatiotemporal dynamics of complex neural processes like voice and language processing.

### Limitations

A major limitation in our study is overall heterogeneity in methodology. As mentioned in our discussion of results, differences in the techniques used to identify VOICE+ and STIM+ channels may not reveal the full extent of functional versatility in these regions. In particular, our investigation likely revealed both essential and ancillary voice-encoding regions but only those regions that were essential for language. Further, the wide variety of tasks and behaviors classified as “disturbance” during clinical stimulation mapping procedures makes it difficult to identify exactly which processes were disrupted and, therefore, to attribute any precise function to the regions identified (see Borchers et al., 2012 for a review).

This examination was additionally constrained by clinical utility, both in terms of the distribution of sEEG electrodes implanted and the channels stimulated in each patient. It is therefore likely that the networks subserving voice encoding and language are more extensive than observed in this study.

### Conclusions and Future Directions

In this exploratory analysis, we identified regions of the left posterior STG/STS that exhibit functional heterogeneity in voice encoding and language. These findings indicate that some regions traditionally labeled as “temporal voice areas” (TVAs) are not solely dedicated to the encoding of human voice and may participate in a diverse range of neural processes.

Furthermore, we demonstrate that these auditory association areas somehow play a critical role in the production of language and that the strength of voice encoding is not significantly weaker in regions with overlapping function compared to voice-encoding regions in the anterior STG/STS that were not identified as critical for language.

Given the relatively narrow range of tasks employed during stimulation mapping procedures, it is not possible to conclude that the channels labeled strictly as VOICE+ in this study are purely voice encoding regions. Prior research suggests that the anterior STG/STS is implicated in phoneme-, word-, and sentence-level speech comprehension (DeWitt & Rauschecker, 2012; Friederici et al., 2003; Humphries et al., 2001; Obleser et al., 2006), as well as more generally in auditory categorization (Leaver & Rauschecker, 2010; Liebenthal et al., 2010). It is therefore possible that functional heterogeneity within classically defined “voice areas” is even greater than estimated in this study. A full account of functional diversity within voice-encoding areas may thus require a wider variety of tasks. Future research that examines the functional roles of these areas throughout the STG/STS in a more granular fashion may not only strengthen observations of versatility, but also improve our understanding of how discrete regions within the voice-encoding network may differentially contribute computationally to auditory processing.

Investigation of voice processing has traditionally relied on neuroimaging and/or electrophysiological recordings to identify areas which show a preference for human voice over other categories of auditory stimuli. As discussed, these techniques provide the spatial and temporal resolution necessary to map the overall network of regions contributing to voice selectivity; however, they do not have the specificity to identify which of these areas may be critical to voice encoding. Future studies may be able to differentiate crucial versus participatory areas by including cortical or transcranial stimulation in auditory categorization tasks. There is a paucity of stimulation studies investigating the role of voice-selective sites in auditory categorization, though Bestelmeyer et al. (2011) did use transcranial magnetic stimulation to show that TVAs in the right hemisphere are particularly important for voice/non-voice discrimination. Using a combination of techniques with high sensitivity and specificity may help in elucidating the regions and neural computations underlying voice processing, including higher order representations such as identity and talker characteristics.

